# Fundamental Theory of the Evolution Force (FTEF): Gene Engineering utilizing Synthetic Evolution Artificial Intelligence (SYN-AI)

**DOI:** 10.1101/585042

**Authors:** L.K. Davis, R.M. Uppu

## Abstract

The effects of the evolution force are observable in nature at all structural levels ranging from small molecular systems to conversely enormous biospheric systems. However, the evolution force and work associated with formation of biological structures has yet to be described mathematically or theoretically. In addressing the conundrum, we consider evolution from a unique perspective and in doing so we introduce “*The fundamental Theory of the Evolution Force*”. Whereby, the driving force of evolution is defined as a compulsion acting at the matter-energy interface that accomplishes genetic diversity while simultaneously conserving structure and function. By proof of concept, we attempt herein to characterize evolution force associated with genomic building block (GBB) formation utilizing synthetic evolution artificial intelligence (SYN-AI). As part of our methodology, we transform the DNA code into time dependent DNA codes based upon DNA hierarchical structural levels that allow evolution by the exchange of genetic information like the swapping of building blocks in a game of Legos. Notably, we not only theorize and mathematical describe the evolution force herein, but have written a set of 14-3-3 docking genes from scratch.

## 1. Introduction

Synthetic evolution artificial intelligence (SYN-AI) is an evolution based AI that writes genes from scratch by reconstructing the evolution force. The evolution force may be described as a compulsion acting at the matter-energy interface that drives molecular diversity while simultaneously promoting conservation of structure and function. The effects of the evolution force are manifested at all levels of life and is responsible for such processes as the formation of genes and gene networks. Herein, we introduce the “*Fundamental Theory of the Evolution Force (FTEF)*” and attempt to theorize and mathematically describe formation of genomic building blocks (GBBs) as well as write genes from scratch utilizing *FTEF*. From our perspective genomic building blocks are short highly conserved sequences formed as evolution artifacts. It is not difficult to assert that DNA and protein are matter based computer programs. When viewing genes from the perspective of a computer algorithm, genomic building blocks are analogous to fundamental programming blocks. SYN-AI identifies evolution force promoting formation of these programming blocks and applies natural selection algorithms to locate evolutionarily fit DNA crossovers.

In constructing our model, consideration of evolution force was limited to what we believe to be the four most fundamental molecular based evolution engines, (i) evolution conservation, (ii) wobble, (iii) DNA binding state and (iv) periodicity. Whereby, strong association between evolutional conservation of DNA/protein sequence and effects on function has long been recognized by molecular biologist (1–5).

Wobble is classically defined as genetic diversity within the third codon with conservation of amino acid sequence (6–12). We expand wobble’s definition to the achievement of genetic diversity with simultaneous conservation of structure, thusly allowing wobble to be quantifiable at all structural levels. We establish DNA binding states as evolution engines based on the assumption that the association of energy and life is inseparable and assert that interaction of evolution force at the matter-energy interface may be characterized by DNA binding states (13–16). A strong correlation between sequence periodicity and conservation of structure and function has been demonstrated by Fourier spectrums (17–19). Wherein, it is widely accepted that nature conserves structures that contribute to survival of an organism. Thusly, we assume that such sequences must occur at higher periodicity.

When applying *FTEF,* SYN-AI integrates a gene-partitioning model that assumes contemporary genes evolved from a single ancestor that expanded to the modern gene pool. Thusly, *FTEF* is in agreement with the “Universal Ancestor” and LUCA “Last Universal Common Ancestor” models, (20, 21). Gene partitioning allows transformation of gene sequences to DNA secondary (DSEC) and DNA tertiary (DTER) codes characterized by DNA hierarchical structure levels. The aforementioned introduce a time dimension to the DNA code allowing fast-forwarding of the evolution process, while dampening the effects of point mutations that lead to disruption of protein function. Application of time based DNA codes also allows for conservation of global and local protein architecture. The DSEC describes evolution on the scale of genomic building block formation, wherein the DTER scales evolution an order of magnitude higher to the formation of super secondary structures. Thusly, exchange of genetic information during synthetic evolution is like the swapping of genomic building blocks in a game of Legos and is in agreement with the ‘Domain Lego’ principle described in (22, 23).

Herein, SYN-AI was utilized to write a set of 14-3-3 docking genes from scratch. According to the “*FTEF*”, we identified genomic building block formations across single and multi-dimension planes of evolution. The *Bos taurus* 14-3-3 ζ docking gene was transformed into a time-based DSEC and genomic building blocks identified by evolution force analysis. Subsequently, synthetic super secondary structures were engineered by random selection and ligation of genomic building blocks. Selection was limited by application of natural selection protocols using Blosum 80 mutation frequency and PSIPRED 4.0 secondary structure based algorithms. Utilizing the aforementioned approach, SYN-AI constructed a library of 10 million genes that was reduced to three architecturally conserved 14-3-3 docking genes by subsequent rounds of selection, whereby synthetic and parental protein active sites were overlapped for theoretical closeness. Notably, we present evidence that SYN-AI was able to write functional 14-3-3 ζ docking genes from scratch and present the first theorization and mathematical modelling of the evolution force.

## 2. Theory

### 2.1 Fundamental Theory of the Evolution Force

We state herein that the evolution force is a compulsion acting at the matter-energy interface that drives genetic diversity while simultaneously conserving biological structure and that the dynamics of the matter-energy interface do not act independently of evolution’s tendency toward conservation. We hypothesis that the four fundamental engines driving gene evolution are (1) evolution conservation, (2) wobble, (3) DNA binding state and (4) periodicity. Whereby, evolution force generated about these engines may be solved for according to the postulates stated below:

***Postulate 1*** - A natural selection system will generate sequences exhibiting positive variation from the mean of a population of randomly evolved sequences occurring during an evolution instance. Whereby, such sequences will exhibit greater evolutionary conservation of the parental sequence.
***Postulate 2*** - Due to degeneracy of the genetic code as described in (Crick, 1966), a natural selection system will generate sequences that exhibit higher conservation of protein structure than expected based on mean DNA similarity. The aforementioned is characterized as wobble and considered an artifact of the evolution force.
***Postulate 3*** - Evolution force regulates molecular diversity at the matter-energy interface in the form of Gibb’s free energy dependent DNA base stacking interactions. Thusly, evolution force may be characterized by DNA binding states.
***Postulate 4*** - Evolution has a tendency to repeat structures that contribute to survival of an organism, wherein structures that contribute to function occur more frequently. Thusly, evolution force may be solved as a function of sequence periodicity.

### 2.2 Four Fundamental Evolution Engines

#### 1) Evolutionary Sequence Conservation

Sequence conservation is strongly correlated with ligand binding, structure of active sites, protein-protein interaction (PPI) interfaces and functional specificity (1). In a study of DNA binding proteins, it was demonstrated that functionally essential residues are more highly conserved than their counterparts (24). It has also been established that evolutionary conservation of amino acids contributes to the protein stable core (25). *FTEF* describes genomic building block formation as a function of evolutionary conservation. Whereby, evolution force about evolution conservation engine *ϵ* describes conservation at DNA and protein levels and is a function of evolution vectors 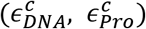. These position vectors reflect DNA crossover homology to the parent sequence in respect to a rigid body comprising the full enumeration of DNA crossovers occurring during an evolution instance. Thusly, they characterize phylogenetic history of a sequence. They are functions of DNA and protein similarity vectors (*X*_*i*_, *X*_*j*_) that compare recombinant sequences to parental in terms of physiochemical properties, volume, hydrophobicity and charge. Whereby, relative position of DNA crossovers to the rigid body describes evolutional advantageousness of a sequence and is solved by weighting similarity vectors by evolutional weights (*W*_*d*_, *W*_*p*_).

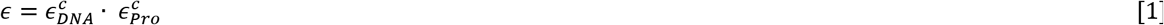

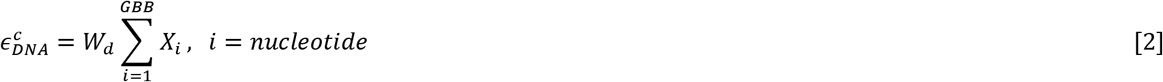

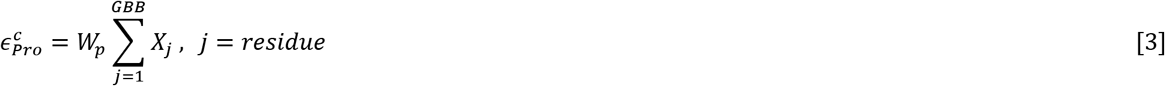

Evolution weights *W*_*d*_ and *W*_*p*_ are measures of central tendency that describe the rigid body center of gravity and are functions of recombinant pool mean DNA 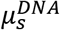 and protein 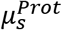 similarity. Thusly, they describe positional vectors of all DNA crossovers occurring during an evolution cycle. Mean similarity is solved by the summation of DNA and protein similarity vectors (*X*_*i*_, *X*_*j*_) occurring within sequence space (*sspace*^*r*^). Where, *sspace*^*r*^ comprises all orthologue and paralogue gene sequences at a selected identity threshold. Evolutional weight is solved in respect to the total number of DNA crossovers (N), thusly reflects full enumeration of DNA crossovers occurring within the evolution potential field and scales the DNA crossover instance in comparison to all crossovers occurring during phylogenetic history of the gene sequence space.

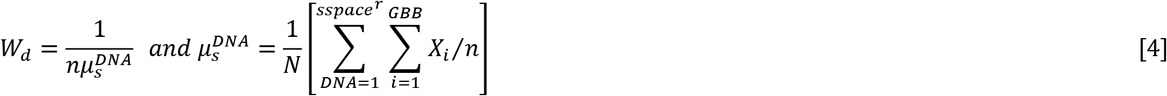

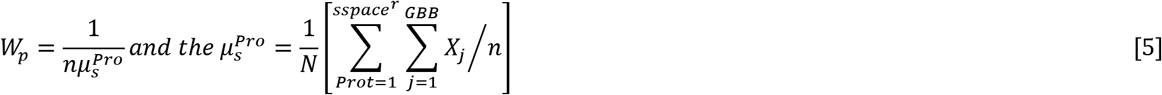

#### 2) Molecular Wobble

Wobble is a result of the genetic code’s evolution, whereby Crick (45) hypothesized evolution from a simple triplet code expressing a few amino acids. A later study by Wu suggests evolution of the modern code from an intermediate doublet system, wherein only the first and second codon positions were read (26). These hypotheses are corroborated by evolution remnants displayed in aminoacyl tRNA synthetases (27). *FTEF* views wobble as an evolution artifact and an indicator of natural selection. Whereby, wobble is solved by overlapping evolution position vectors 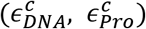. The relationship is reflective of hierarchical sequence transitions across the same sequence. *FTEF* designates wobble *ω*_*m*_ as a function of genetic displacement *x* over time, where genetic displacement of the protein position vector respective to the DNA position vector is described by 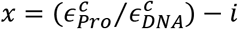. Identity vector *i* = 1 is the expected position of the DNA crossover in evolution space and reflects the rigid body center of mass. Time *t* is the number of evolution cycles.

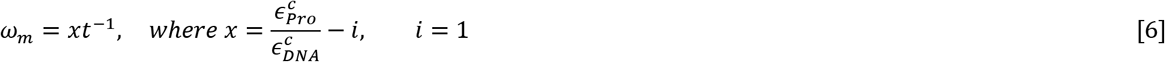

#### 3) DNA Binding States

*FTEF* assumes synthetic evolution processes mimic evolution, thusly DNA binding states occurring during synthetic evolution are analogous to DNA crossovers occurring during meiosis. Current theory suggests that genetic diversity occurs by DNA crossovers and translocations that result in processes such as gene duplication, inversion, insertion and deletion (28, 29). It is also widely accepted that these processes result in relaxation of evolutional stringency, thusly allowing speciation and random point mutations by neutral evolution (30, 31). *FTEF* solves DNA binding states *p*^*i*^ as a function of annealing probability 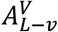, described in (15) and DNA binding probability 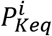 (14). Thusly, *p*^*i*^ is a function of volume exclusion effects and DNA crossover thermodynamic signatures.

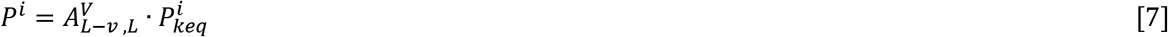

Annealing probability 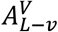 distributes volume exclusion *V*^*α*^ (32) characterizing a DNA hybridization over that of the recombinant pool. *V* defines overlap length characterizing a DNA crossover and *L* defines sequence length. Volume exclusion is a function of the length to volume relationship occurring at the DNA crossover junction, whereby the probability of hybridization decreases beyond a critical volume of the hybridization bubble.

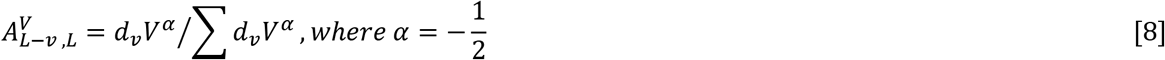

Thermodynamic contributions to the DNA binding state are solved as a function of the equilibrium constant *k*_*eq*_. Where, *k*_*eq*_ of a DNA crossover is an exponential function of Gibb’s free energy. Gibb’s free energy of hybridization is solved by summation of *G*° (*i*) defining standard free energy changes for the 10 possible Watson-Crick nearest neighbors occurring in a DNA crossover (43). Counterion condensation is accounted for by free energy of initiations *G*° (*init w/term G · C*) and *G*° (*init w/term A · T*). Likewise, *G*° (*i*) incorporates an entropic penalty *G*°(*sym*) for maintaining C2 symmetry of self-complimentary sequences.

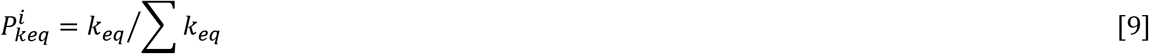

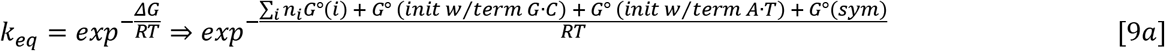

#### 4) Periodicity

A strong peak at frequency 1/3 is observed in Fourier spectrums of genome coding regions (33), suggesting the presence of selection pressure. Three base periodicity also allows characterization of species based upon Fourier spectrum (34). *FTEF* characterizes periodicity *P*^*π*^ as the distribution of genomic building block frequency *f*_*ij*_ over global frequency Z. *f*_*ij*_ describes oligonucleotide *i* and peptide *j* sequence homolog occurrences within the target gene, where Z is the summation of all DNA crossover frequency within the orthologue sequence space, thusly, describes periodicity across the phylogenetic history of the gene. *P*^*π*^ compares selectivity of a DNA crossover to adjacent sequences at DNA and protein levels. Whereby, sequences displaying high periodicity are reflective of selection pressure by the evolution force.

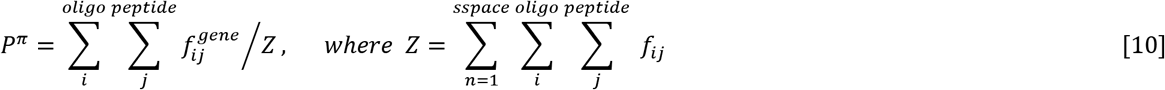

### 2.3. Analyzing Evolution Force utilizing the Linear Model

The *“Linear Model”* considers evolution force at the DNA and protein level ignoring transitory effects on mRNA transcripts. The *FTEF* predicts genomic building block formation by identifying evolution force characterizing DNA crossovers. Sequences are viewed as particles having momentum through an evolution potential field generated by a rigid body comprised of DNA crossovers occurring during the evolution process back to the LUCA. We apply Newton’s second law of motion to describe particle momentum across the evolution potential field as described by *p* = *mv* ⇒ *ϵ* · *ω*_*m*_. *FTEF* captures a snapshot of evolution by setting particle mass analogous to evolution engine *ϵ* and genetic velocity analogous to wobble *ω*_*m*_. Thusly, evolutional momentum *p* reflects change in sequence homology during evolution and the rate of mutation. Acceleration thru the potential field is the derivative of mutation rate *ω*_*m*_. Mutation rate describes evolution of the gene as well as captures some remnants of the evolution of the genetic code.

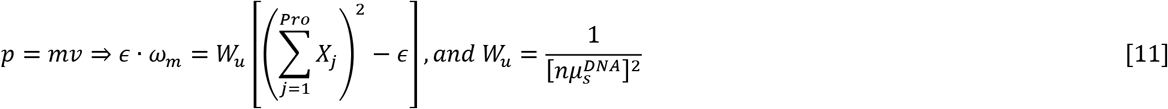

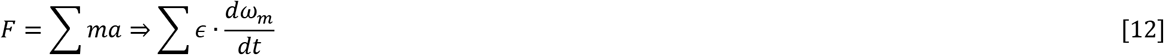

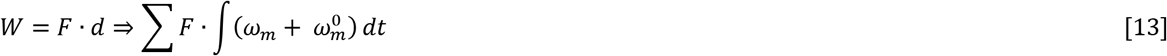

#### 2.3.1 Analyzing Evolution Dynamics Utilizing the Linear Model

In order to elucidate evolution dynamics, *FTEF* must describe the relative position of the parent sequence to the rigid body. As the initial position of the parent sequence within the evolution potential field cannot be ascertained. We show that momentum p of the parental sequence is dependent on evolution vectors 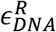 and 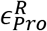 describing relative position of parental DNA and protein sequences in evolution space. The aforementioned are a function of the summation of identity vectors characterizing an ideal DNA crossover of 100 percent sequence homology as well as weighted by the recombinant pool. Thusly, *FTEF* views the parent sequence analogous to an ideal DNA crossover, wherein its position is determined relative to the rigid body in evolution space.

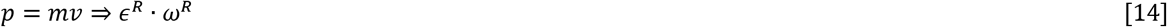

*FTEF* solves *PE* as a steady state where evolution potential is a function of potential mass *m*_*φ*_ and genetic distance *h*. Potential mass *m*_*φ*_ = *ϵ*^*R*^ − *ϵ* reflects differential homology between recombinant and parental sequences. Displacement *h* describes genetic distance of the DNA crossover to the parent sequence within the evolution potential field, idyllic DNA crossovers are characterized by smaller *h* values. The system is pliable as the evolution potential field changes with each DNA crossover instance generating fluctuations in sequence homology. Sequences characterized by high PE display low evolvability. The system’s kinetic energy reflects magnitude of evolution force applied on the sequence space and is solved as a function of the evolution conservation engine and mutation rate (wobble). Alternatively, KE is solved as a function of acceleration *a*_∈_ and distance vector *x*.

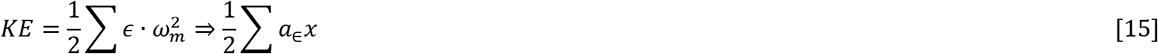

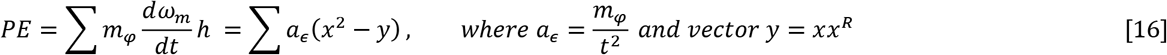

The relationship between wobble and incremental potential energy changes within a sequence space during an evolution instance is described by a first order differential equation *dPE* = 2*a*_*∈*_*xdx*. The potential energy vector allows comparison of sequence spaces in respect to mutation rate, where low values are indicative of greater evolvability.

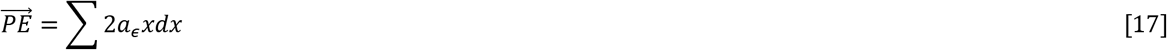

Total energy reflects evolutional advantageousness of the system.

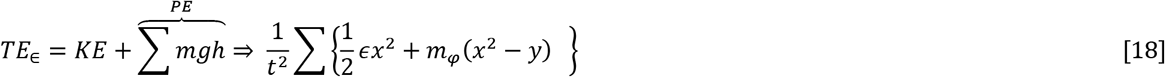

The relationship between wobble and TE is a function of the acceleration occurring about the wobble engine and the resultant of the evolution conservation *ϵ* engine and the potential mass *m*_*φ*_ vector. When evaluating the TE vector with respect to time, change in 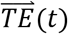 is a function of jerk about the wobble engine. Thusly, reflects mutation rate dynamics of the recombinant pool respective to the phylogenetic history of the gene.

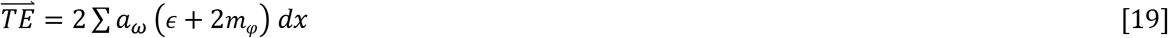

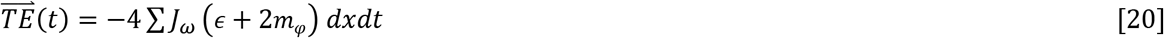

The Lagrangian 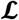 of the system describes the path of the least evolutional resistance. The optimal path for gene formation is enumerated by summation of 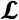 characterizing DNA crossovers occurring within each genomic alphabet of the DESC over the evolution period. The state 𝒮 describes the evolutional equilibrium. Less negative states indicate highly evolvable sequence spaces.

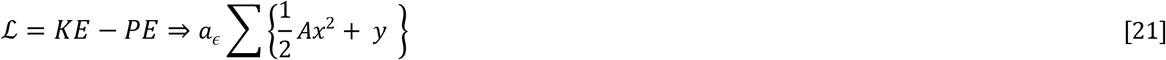

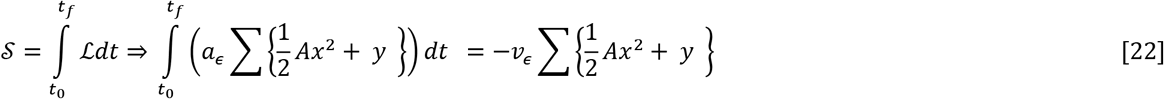

### 2.4 Analyzing Evolution Force utilizing the Rotation Model

The ‘Rotation Model’ analyzes evolution force associated with genomic building block formation as a function of evolutional inertia. Whereby, DNA crossover instances are analogous to particles revolving a rigid body of particles formed by the recombinant pool. Evolution force is a function of moments of inertia characterizing the DNA crossover and its acceleration about the respective evolution engine. Evolutional inertia of the particle is described by *I* = *ϵr*^2^. Where, radius *r* describes standard deviation of evolution position vector *ϵ* from the rigid body. Thereby, evolution force *τ*_*ϵ*_ is solved as a measure of central tendency respective to the phylogenetic history of the gene.

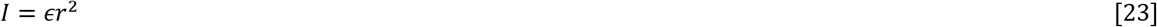

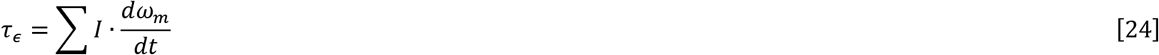

Work applied on the system is a function of evolutional torque and genetic distance *θ*. Where, 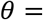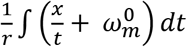 describes the genetic step in respect to the rigid body.

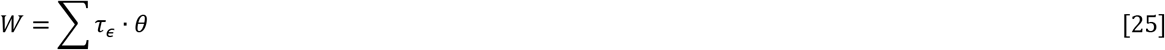

The system’s kinetic energy is a function of inertia about evolution engine E and velocity of the particle across the potential field.

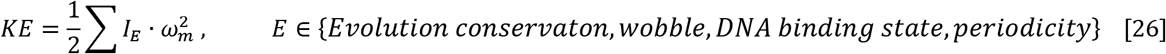

TE is the sum of inertial kinetic and inertial potential energies. Where, evolution potential energy PE is a function of inertial vector *I*_*φ*_ characterizing potential moments of inertia about the evolution engine. The state *S* is characterized by the difference in DNA crossover angular momentum *L*_*KE*_ in direction of the kinetic energy vector and its angular momentum *L*_*PE*_ in the direction of the potential energy vector.

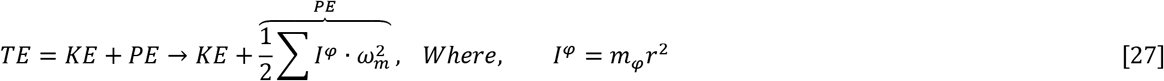

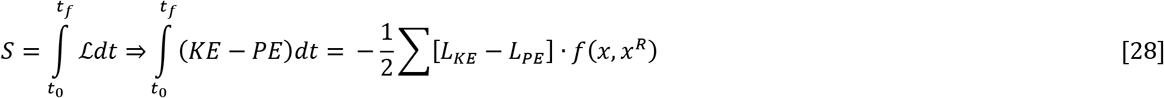

Variation in the system’s total energy in respect to wobble is a function of torque about the evolution engine and incremental change in the wobble distance vector (*dx*). Whereby, instantaneous change in the system’s state is a function of the angular momentum about the evolution engine and (*dx*).

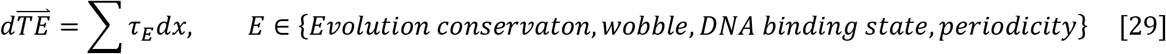

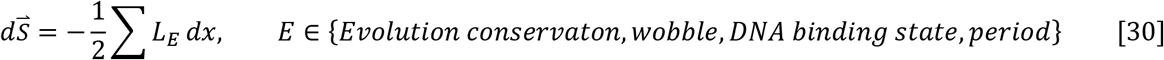

## 3. Evaluating Synthetic Structures

*FTEF* defines wobble as the conservation of structure in face of genetic diversity. When wobble occurs at the macroscopic level and higher, the tendency is referred to as structural wobble. An example of structural wobble is phyllotaxis, the arrangement of leaves on plants and deformation configurations seen on plant surfaces described in (42). Wherein, it is explained that the aforementioned may be understood as energy-minimizing buckling patterns of a compressed shell on an elastic foundation. These Fibonacci-like patterns occur across plant species encompassing a broad range of genetic diversity. The *FTEF* solves for structural wobble as a conditional probability of target structure similarity to the native state. The probability that a state *x*_*s*_ formed during synthetic evolution will share homology with the native state is based upon closeness probability *θ*_*i*_, where *i* is an element of set *P* comprising physiochemical properties volume, hydrophobicity and folding propensity.

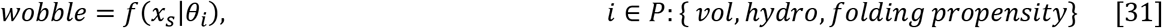

To prevent protein structural perturbations, SYN-AI performs high-resolution pattern recognition by analyzing discrete sequence spaces occurring across protein structures. SYN-AI walks the genomic building block protein sequence in steps of three residues, where propensity of characteristic (*i*) within the sequence space is summated as illustrated in **Eq. 32**. Structural propensity (*p*) within discrete sequence spaces is fingerprinted by probability density function (δ) as illustrated in **Eq. 33**. Whereby, area under the density curve ∫ *P dp* is normalized by partition function *σ* characterizing summation of characteristic *i* occurring across the structure. The aforementioned procedure allows SYN-AI to characterize the taste of the sequence space. Whereby, proteins may be characterized by diverse flavors describing discrete changes in residue volume, hydrophobicity and folding propensity occurring both locally and globally. Theoretical closeness is described by probability *θ*_*i*_ and solved as a function of synthetic and native states, **Eq. 34**.

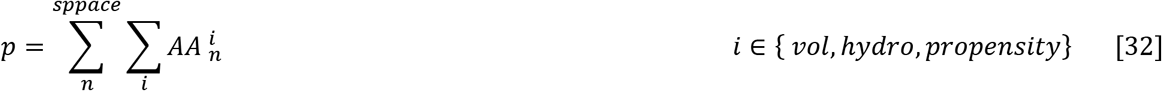

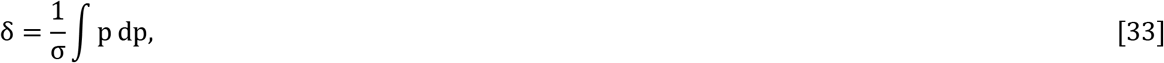

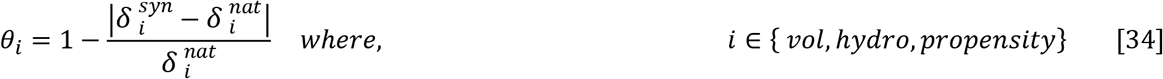

In solving the probability of structural state *x*_*s*_, *θ*_*i*_ is factored across *n* sequence spaces comprising the structure. Where, *i* is an element of secondary, super secondary and quaternary structural groups.

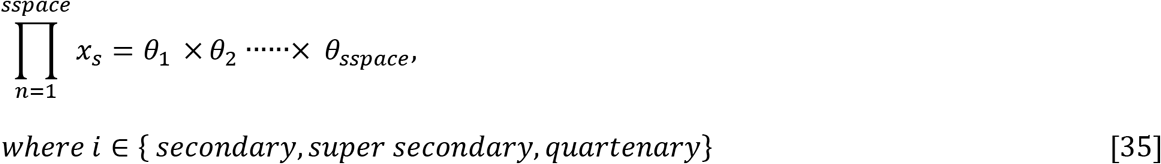

*FTEF* solves wobble as a function of average closeness 〈*Closeness*〉 of synthetic and native states. Whereby, closeness *θ*_*i*_ is summated over *n* discrete sequence spaces comprising the structure and over characteristic (*i*). Where, N reflects the total number of measurements and *i* is an element of physiochemical properties volume, hydrophobicity and folding propensities. Wobble occurring across the structure is solved as a function of 〈*Closeness*〉 and protein similarity *Prot*_*s*_. Thusly, comparing structural conservation to genetic diversity occurring at the protein level.

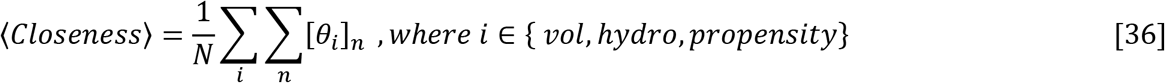

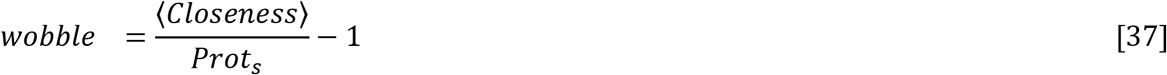

## 4. Methods

### 4.1 High Performance Computing

SYN-AI was performed utilizing the Stampede 2 supercomputer located at the Texas Advanced Computing Center, University of Texas, Austin, Texas. Experiments were performed in the normal mode utilizing SKX compute nodes comprising 48 cores on two sockets with a processor base frequency of 2.10 GHz and a max turbo frequency of 3.70 GHz. Each SKX node comprises 192 GB RAM at 2.67 GHz with 32 KB L1 data cache per core, 1 MB L2 per core and 33 MB L3 per socket. Each socket can cache up to 57 MB with local storage of 144 /tmp partition on a 200 GB SSD.

## 5. Results and Discussion

### 5.1 Analysis of Evolution Force

In order to simulate evolution, we partitioned the parental *Bos Taurus* 14-3-3 ζ docking gene in DNA secondary (DESC) and tertiary codes (DTER) based upon DNA hierarchical structural levels in agreement to the “*Domain Lego*” theory (22, 23). SYN-AI simulated DNA crossovers and identified genomic building blocks by performing DNA hybridizations within the DESC. Whereby, *FTEF* assumes that modern genes have a common ancestor that partitioned over time via DNA crossovers and that genetic diversity occurred by processes such as gene duplication, inversion, insertion and deletion (28, 29). Hybridization partners were randomly selected across an orthologue/paralogue sequence space and evolution force analyzed utilizing “*Linear*” and “*Rotation*” models introduced in the “Theory” section.

**Figure 1.**
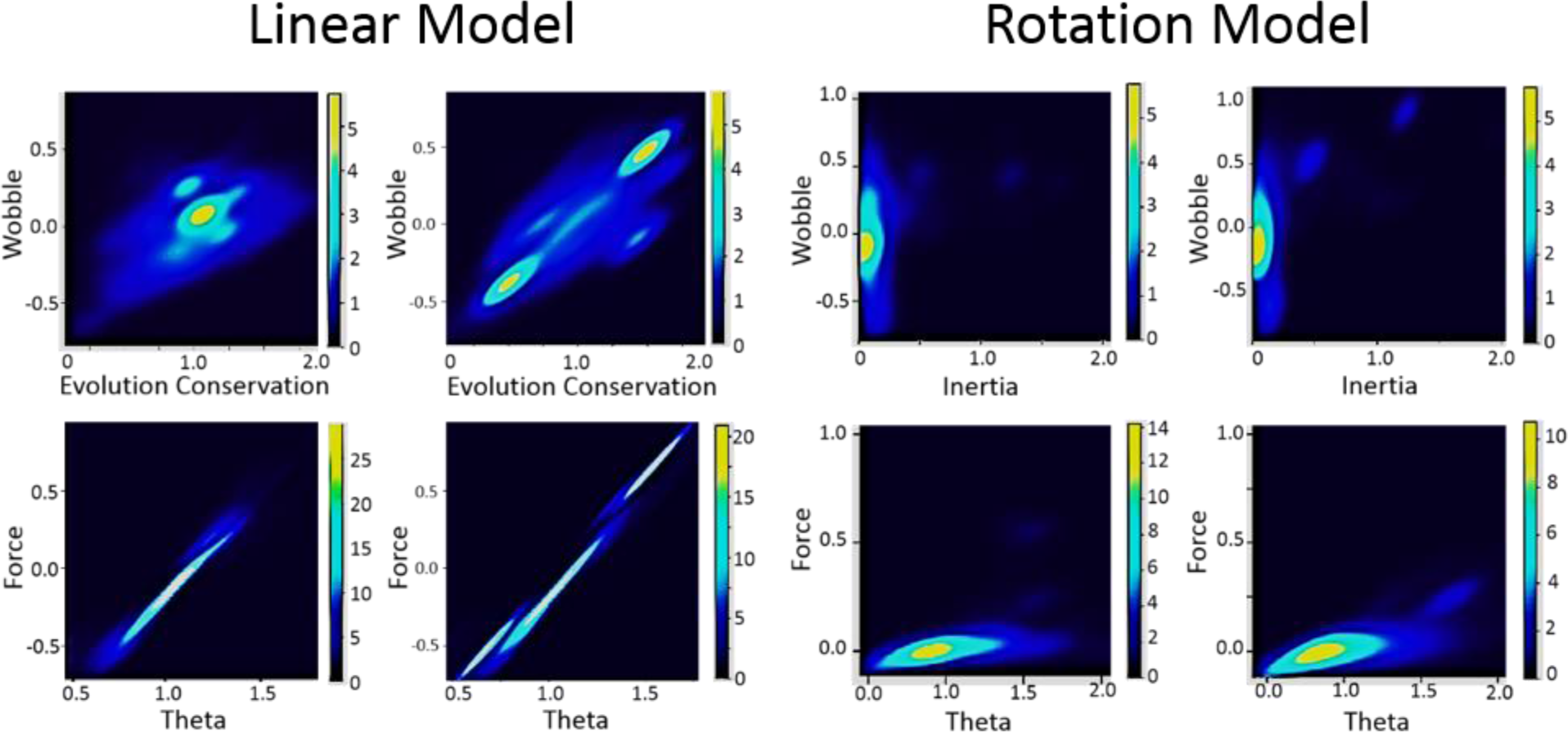
Evolution Force Linear vs. Rotation Model. Evolution force and work was analyzed utilizing the Linear Model as described in the “Theory section”. Evolution force distribution sequence space 1 of the DESC (**Top Left**). Evolution force distribution sequence space 2 (**Top Right**). Work distribution sequence space 1 (**Bottom Left**). Work distribution sequence space 2 (**Bottom Right**). Evolution force and work was analyzed utilizing the Rotation Model. Evolution force distribution sequence space 1 (**Top Left**). Evolution force distribution sequence space 2 (**Top Right**). Work distribution sequence space 1 (**Bottom Left**). Work distribution sequence space 2 (**Bottom Right**).

The Linear Model resulted in broad distribution of evolution force and low resolution of genomic building blocks. However, successfully captures the formation of multiple evolution foci, **Fig. 1** (Linear Model, Top Right). The formation of multiple foci is due to wobble occurring in multiple directions indicating the presence of strong selection and deselection mechanisms. Localization of foci in positive evolution space indicates convergence toward WT and selection pressure based on biological function. Localization of foci in negative evolution space suggests loss of evolutional stringency due to gene duplication. Whereby, the evolution force exerts little effort to maintain integrity of these sequences. Relaxation of evolutional stringency is corroborated by colocalization of foci to a region of low evolutionary conservation indicating occurrence of neutral evolution and point mutations. Deselection was followed by strong selective pressure due to a change in biological function causing formation of foci in negative evolution space. In contrast, the Rotation Model achieved high-resolution of genomic building blocks, **Fig. 1.** Distribution of evolution force occurred near the gravitational epicenter indicating that majority of DNA crossovers were non-informative with a minimum amount of work exerted by the evolution force.

Our experiments corroborate findings of Aravind that suggests there exists a complex interplay between evolution conservation and achievement of genetic diversity (35). In agreement, our experiments demonstrate that the evolution force performed work in the direction of structural conservation and genetic diversity, **Fig. 1** (Bottom). Whereby, work applied in these directions was equally distributed allowing the mechanisms to offset. The evolution force exerted no overall work on orthologue/paralogue sequence space, thusly corroborates the *FTEF* in that evolution is a low energy system. As anticipated, sequence spaces exhibited variant behavior. We hypothesis that sequence specific variations in evolution force distribution were caused by thermodynamic restraints occurring during evolution. Where, loss of evolutionary stringency generated restraints during DNA recombination, allowing sequences retaining high homology to bind more stably as well as display higher magnitude of Gibb’s free energy, while incurring lower thermodynamic penalties. We hypothesize that such free energy partitions guided evolution processes and are intrinsic components of the evolution force.

### 5.2 Analysis of Synthetic Proteins generated utilizing FTEF

SYN-AI constructed a library of 10 million genes that was reduced to three 14-3-3 ζ monomers. Natural selection was simulated by examining theoretical closeness to native states as described in the Theory section with a minimum closeness threshold of > 90 percent identity. Selection of functional proteins was performed by overlapping synthetic and parental active sites with a minimal theoretical closeness threshold of > 90 percent identity. Following natural selection, three-dimensional structure of 14-3-3 ζ monomers was predicted utilizing I-TASSER (36). I-TASSER data validated that *FTEF* achieved conservation of parental 14-3-3 ζ structure accompanied by subtle changes in protein molecular surface. Synthetic proteins comprised well-formed ligand binding pockets characterized by conserved volume, **Fig. 2**. Synthetic 14-3-3 ζ monomers were confirmed with an average confidence score of 1.51, thusly, predicted structures are very reliable. The parental *Bos taurus* 14-3-3 ζ monomer folded at a confidence of 1.55. Synthetic proteins SYN-AI-1 and SYN-AI-3 folded with TM-scores of 0.93±0.06 and estimated RMSDs of 2.5±1.9Å. Where, SYN-AI-2 folded at a TM-score of 0.92 ±0.06 and an estimated RMSD of 2.6±1.9Å. Thusly, corroborating conservation of global protein architecture and demonstrating that synthetic monomers overlap closely with the parental 14-3-3 ζ monomer.

**Figure 2.**
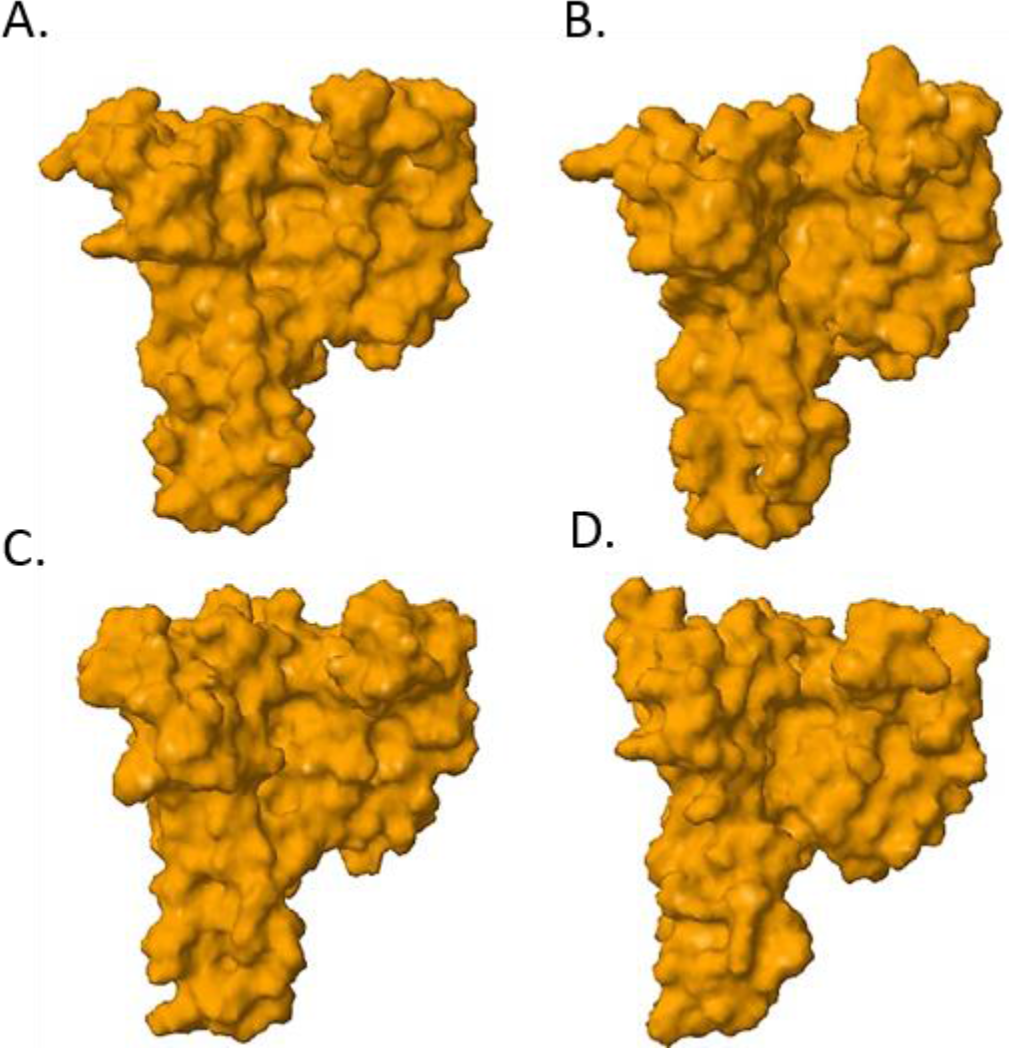
Synthetic 14-3-3 Docking Protein Three-Dimensional Structure. Protein three-dimensional structure was analyzed utilizing the I-TASSER suite (Zhang Laboratory, University of Michigan). Predicted folding of the *Bos taurus* 14-3-3 ζ monomer (**A**), SYN-AI-1 (**B**), SYN-AI-2 (**C**) and SYN-AI-3 (**D**).

In addition to conservation of global and local protein architecture, maintenance of long-range noncovalent interactions is an important detail in protein engineering. As allosteric interactions are important factors in communications between active and ligand-binding sites (37). The effect of synthetic evolution on allosteric interactions was analyzed utilizing the anisotropic network model, ANM2.1 (38). We found that allosteric interactions were conserved in synthetic 14-3-3 ζ monomers with slight modifications in location of nodes and cliques. **Fig.3**. Notably, while cooperative communications were conserved, participating residues were altered. As Clustal Omega sequence alignment and Pylogeny.fr phylogenetic analysis revealed significant sequence divergence between synthetic and parental monomers as characterized by mean sequence identity of 70.25 and mean divergence of 7.33. Close approximation in the location and amplitude of protein nodes despite the significant genetic divergence is corroborated by predicted eigenvectors. Eigenvalues suggests there exists only modest variations in vibrational dynamics of parental and synthetic monomers, **Fig. 4**. Thusly, validating *FTEF* achieves synthetic evolution without disrupting cooperative motions occurring during ligand binding and signal transduction.

**Figure 3.**
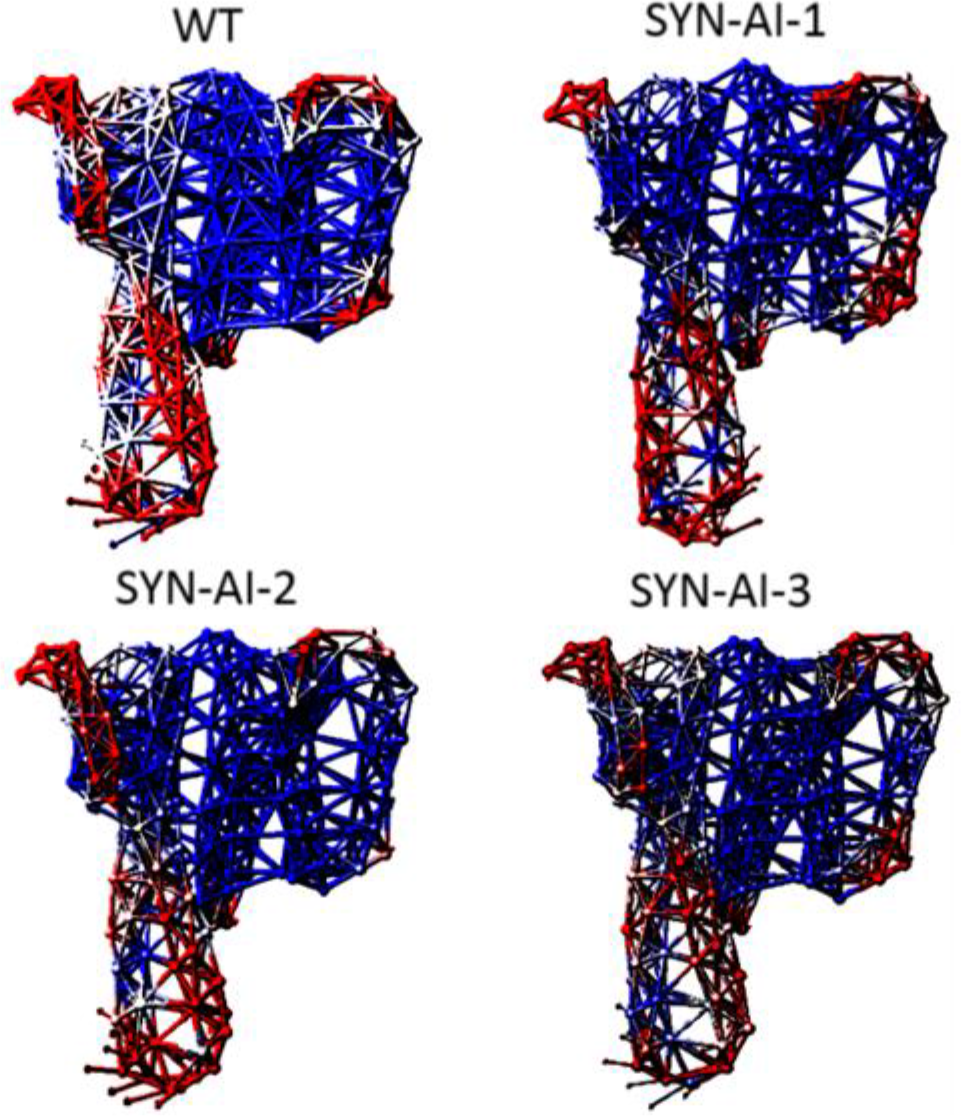
Analysis of Allosteric Interactions. Normal mode analysis was performed utilizing the anisotropic network model ANM2.1, with a Cα cutoff distance of 15 Å and distance weight factor of 0.

Due to conservation of allosteric interactions within the 14-3-3 ζ monomer, it is safe to assume there is no disruption in ligand-binding. Corroborating the aforementioned, vibration potentials of synthetic monomers closely mirrored parental with conservation of the C-terminal hinge bending mechanism, with exception of synthetic protein SYN-AI-1 that displayed vibrational vectors in the opposite direction. Notably, normal mode analysis is in agreement with phylogenetic analysis. As the SYN-AI-1 monomer exhibits greatest divergence from the parental 14-3-3 ζ monomer at 8%, compared to 7 % divergence characteristic of SYN-AI-2 and SYN-AI-3. To corroborate results, mean standard fluctuations were compared to experimental B-factors, **Fig. 5**. There was little variation of ANM2.1 results and experimental B-factors, for parental and synthetic 14-3-3 ζ monomers. Lower estimated experimental peaks near residues 73 and 103 may be explained by bias in temperature factors due to non-natural crystal contacts observed in exposed regions (44). Further, we may have achieved even better agreement by analysis of homodimers as opposed to 14-3-3 ζ monomers. Thusly, we may assume estimation of fluctuation dynamics are accurate.

**Fig. 4.**
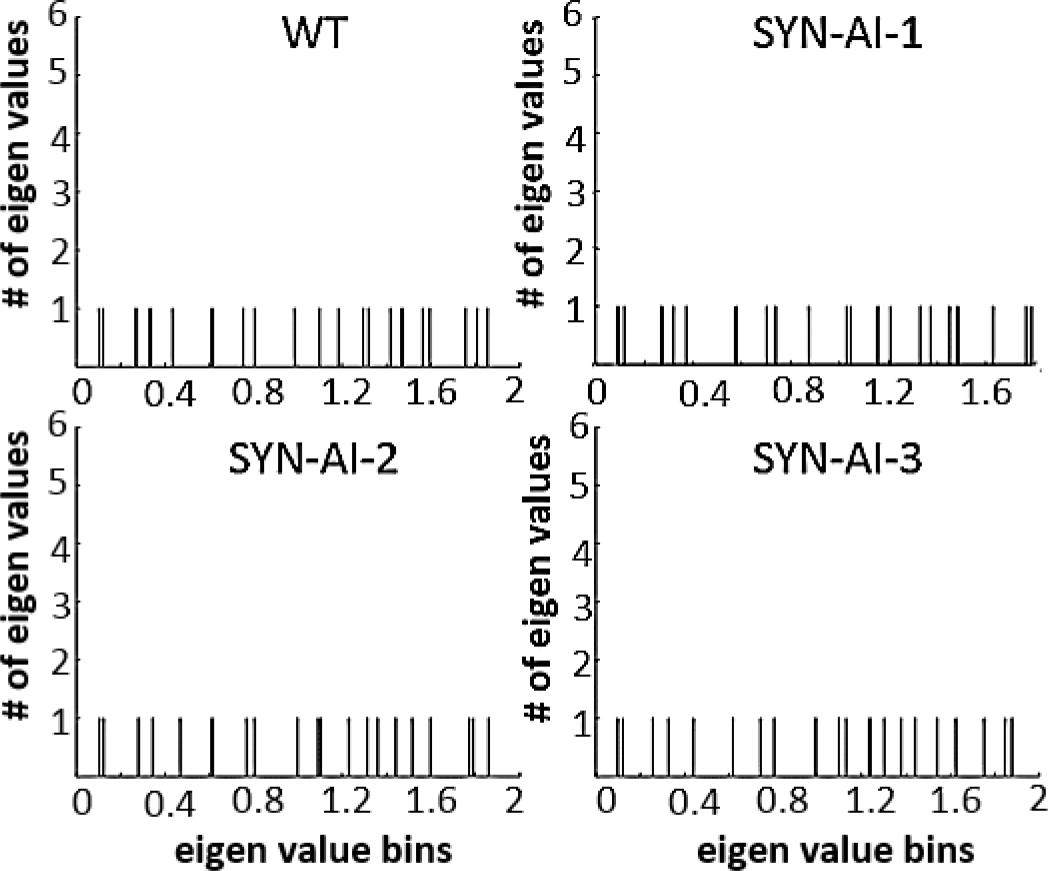
Normal Mode Analysis. Eigenvalues of the *Bos taurus* 14-3-3 ζ monomer and synthetic proteins were calculated utilizing the anisotropic network model.

**Figure 5.**
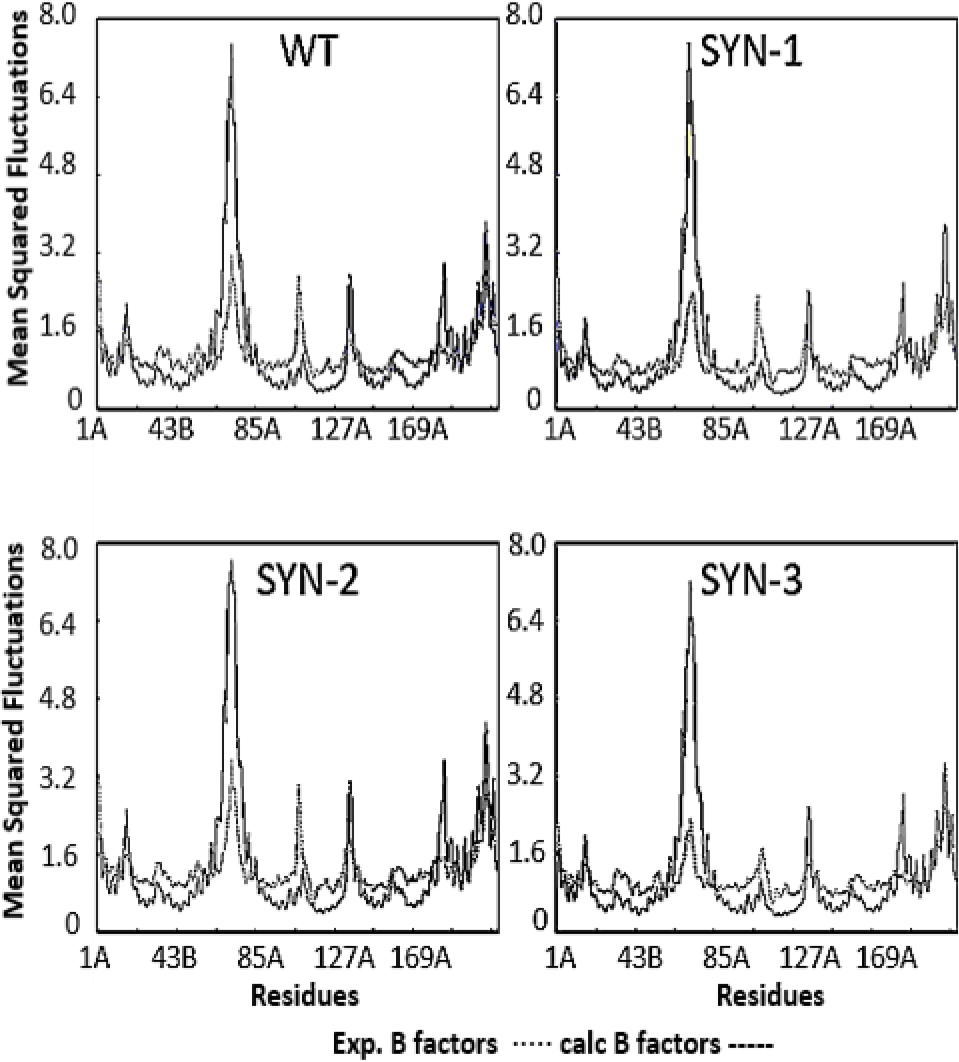
Experimental B-factors. Normal mode analysis of the parental 14-3-3 ζ monomer and synthetic proteins was analyzed utilizing the anisotropic model. Fidelity of normal mode analysis was assessed by comparing predicted mean square fluctuations to experimental B-factors.

When comparing energy deformations occurring within synthetic and parental 14-3-3 ζ monomers, peak pattern is conserved with only slight modifications in peak strength, location and bandwidth, **Fig. 6**. Thusly, potential energy fluctuations occurring within synthetic monomers approximate those of the parental 14-3-3 ζ monomer, **Fig. 6** (A). Occurrence of analogous but slightly modified energy deformation patterns in synthetic 14-3-3 ζ monomers indicates achievement of refined environmental structural changes without disruption of function. Location of energy deformation peaks is indicative to fusicoccin and protein-protein binding sites such as the R18 peptide within the amphipathic groove as corroborated by Coach and Cofactor ligand-binding analysis, **Fig. 6** (B, C). Sequence alignments show that the FC complex binding site of SYN-AI-1 was fully conserved. While, SYN-AI-2 and SYN-AI-3 FC binding sites were characterized by V46 →A46 and N42 →V42 point mutations, respectively. Thereby, energy deformation fluctuations within SYN-AI-1 are attributed to global architectural changes external to the FC complex binding site. Whereby, energy deformation fluctuations in SYN-AI-2 and SYN-AI-3 are due to global and local mutations. Normal mode analysis of exposed residues is not highly reliable (44). Thusly, to corroborate structural predictions we compared solvent accessibilities of parental and synthetic 14-3-3 ζ monomers, **Fig. 6** (D). Solvent accessibilities overlapped parental, thusly corroborating *FTEF* achieved architectural conservation of the parental *Bos taurus* 14-3-3 ζ monomer.

**Figure 6.**
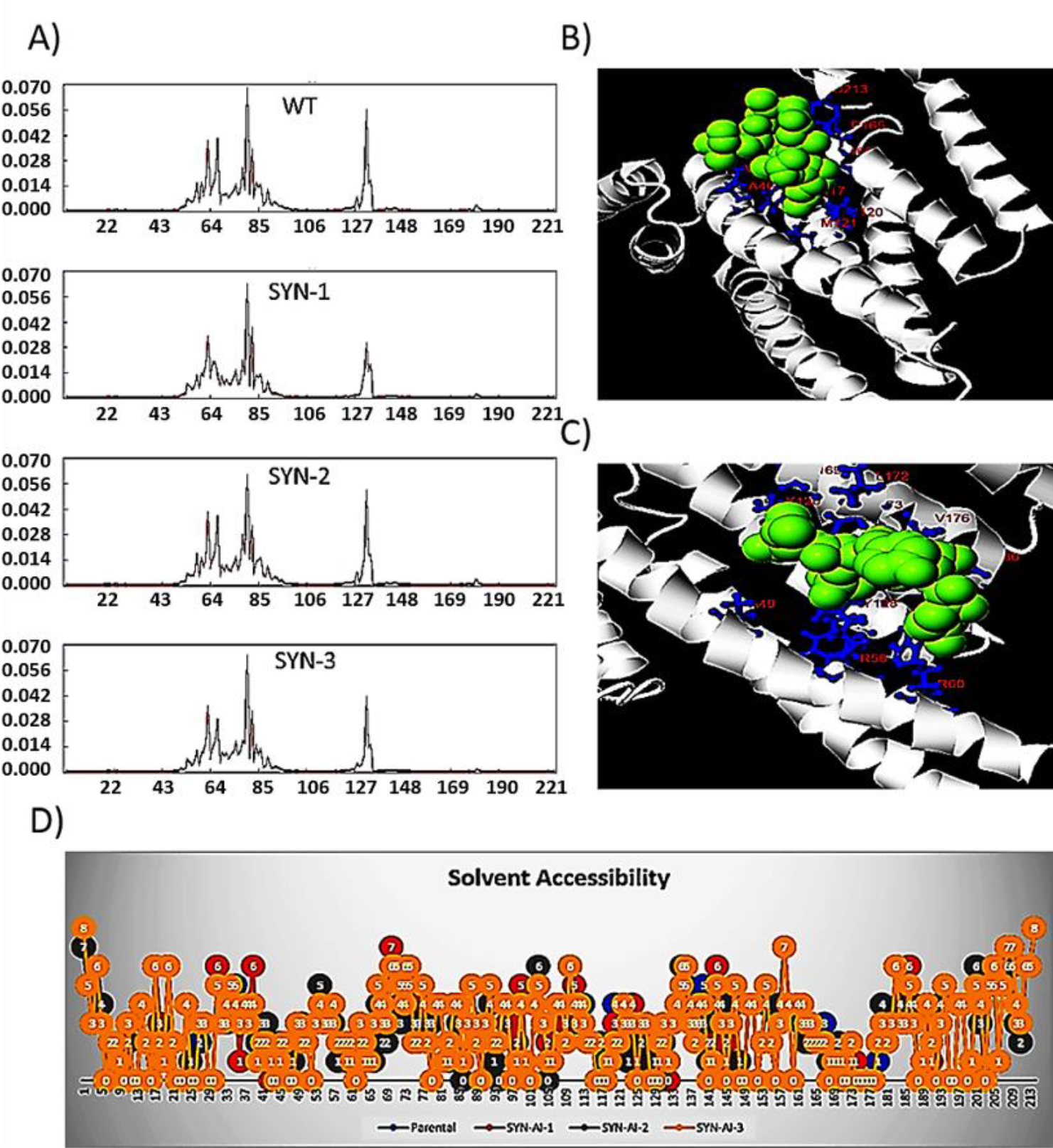
Analysis of Amphipathic Groove. The anisotropic network model was utilized to analyze energy deformations occurring within parental and synthetic 14-3-3 ζ monomers (**A**). Cofactor and Coach were utilized to predict FC complex binding (**B**) and protein-protein interaction (**C**). Solvent accessibility was also analyzed utilizing ANM2.1 (**D**).

Low frequency vibrations occurring within synthetic 14-3-3 ζ monomers were further analyzed utilizing elNemo (39, 40). Utilizing the elastic network model, we demonstrate conservation of the mode 9 “*bend and flex*” mechanism in synthetic monomers, **Fig. 7**. Where, the 14-3-3 ζ monomer closes upon ligand binding locking the substrate into the active site. According to Suhre, amino acid sequences may have evolved so that low-energy barriers are formed when a protein is displaced along normal mode coordinates (39). Notably, our data suggests that *FTEF* successfully captures selective pressures that govern open and closed configurations. It is worth mentioning, while we demonstrated conservation of allosteric interactions within the 14-3-3 ζ monomer, the native *Bos taurus* 14-3-3 ζ docking protein exists as a dimer. Thusly, we analyzed the ability of synthetic 14-3-3 ζ monomers to form homodimers utilizing COTH as described in (41). Saliently, synthetic proteins displayed the potential for dimer formation as illustrated in **Fig. 8**. We also performed normal mode analysis on synthetic 14-3-3 ζ homodimers. Notably, synthetic homodimers conserved the “*bend and flex*” mechanism of the native *Bos taurus* docking protein. Thusly, we demonstrate that *FTEF* not only conserves allosteric interactions in 14-3-3 ζ monomers, but also conserves cooperative communications within synthetic homodimers.

**Figure 7.**
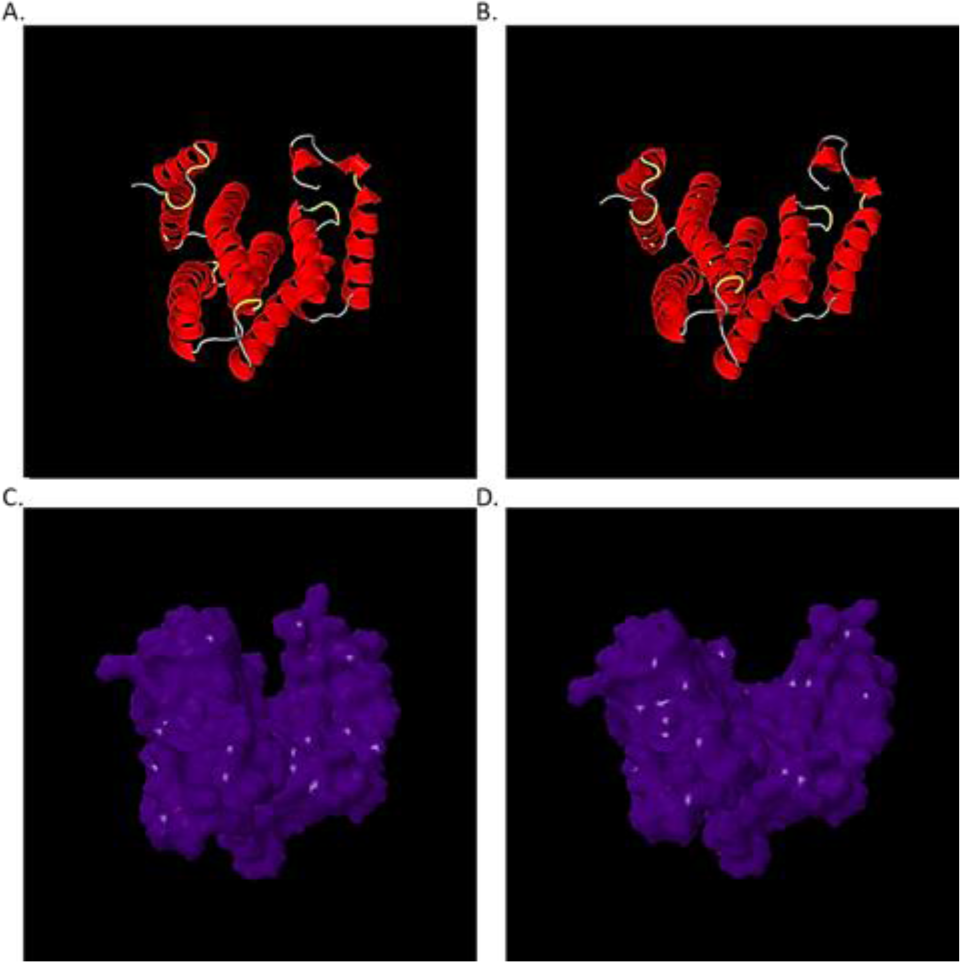
Allosteric Effects of Synthetic Evolution. Normal mode analysis of SYN-AI-3 was performed utilizing the elastic network model, elNemo, with min and max DQ amplitude perturbation of 100. Mode 9 “Bend and flex” mechanism closed configuration (**A, C**), open configuration (**B, D**).

**Figure 8.**
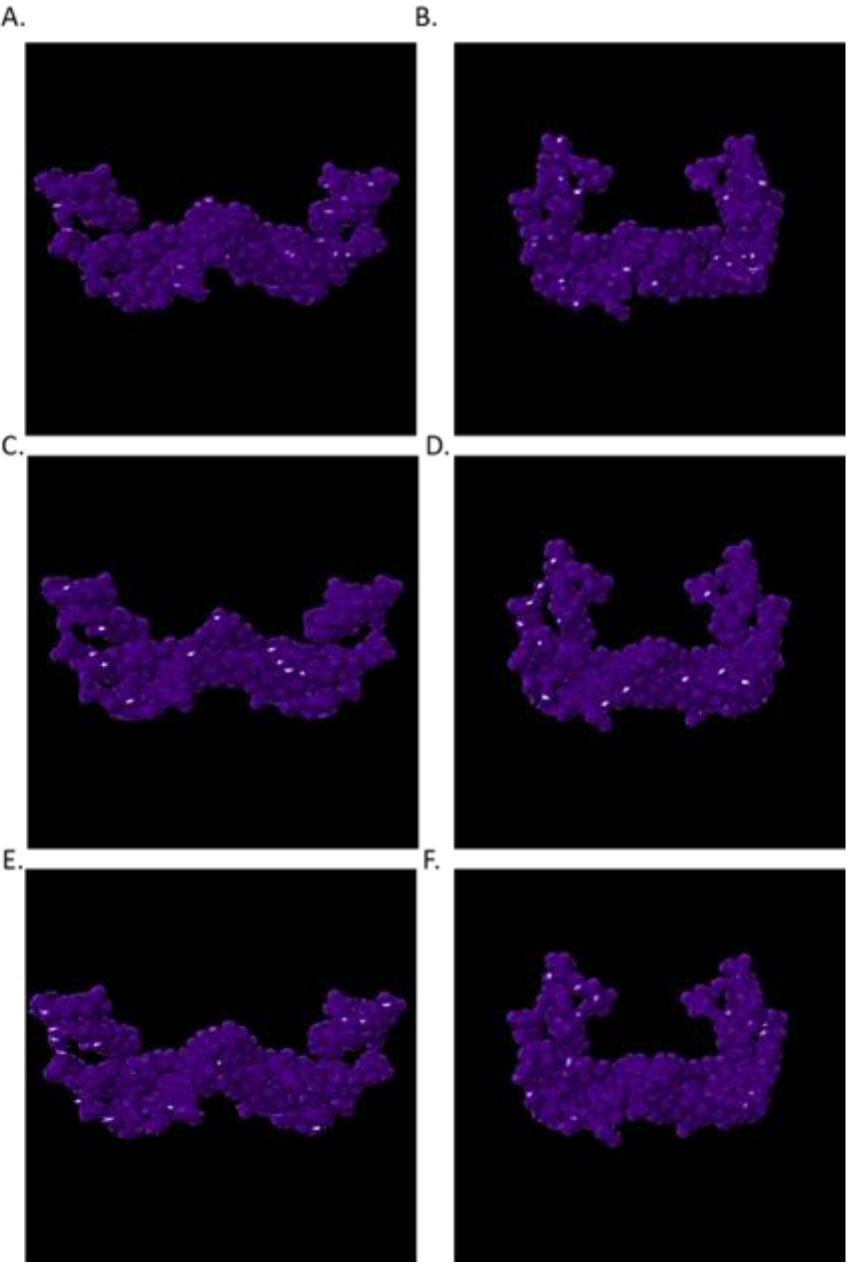
Synthetic 14-3-3 Dimer Formation. Homodimer formation was predicted utilizing COTH. SYN-AI-1 dimer open configuration (**A**). SYN-AI-1 dimer closed config. (**B**). SYN-AI-2 dimer open config. (**C**). SYN-AI-2 dimer closed config. (**D**). SYN-AI-3 dimer open config. (**E**). SYN-AI-3 dimer closed config. (**F**).

## 6. Conclusion

In the current study, we validated ***“** The Fundamental Theory of the Evolution Force: FTEF**”*** by proof of concept. Whereby, synthetic evolution artificial intelligence (SYN-AI) was utilized to engineer a set of 14-3-3 docking genes from scratch utilizing a parental *Bos taurus* 14-3-3 ζ monomer as template for time based DNA secondary (DESC) and tertiary (DTER) codes to guide the engineering process. In contrast to rational design approaches, genes were constructed by random assembly of genomic building blocks identified by analysis of evolution force. By simulating natural selection, SYN-AI was able to engineer synthetic 14-3-3 ζ monomers that display significant divergence from the template gene, while conserving global and local protein architecture. Notably, protein-protein interaction and ligand-binding sites were also conserved. We demonstrated that gene engineering utilizing *FTEF* achieved conservation of 14-3-3 ζ allosteric interactions and the potential for dimer formation. Saliently, cooperative communications were conserved in synthetic 14-3-3 ζ monomers and homodimers. We conclude that *FTEF* gene engineering is an excellent approach to generating functional proteins as well as an excellent methodology for engineering cell signal pathways.

## Author Contributions

R.M. supervised and funded initial aspects of the study. L.K. developed FTEF, designed SYN-AI and wrote the article.

## Acknowledgements

The authors acknowledge the Texas Advanced Computing Center (TACC) at The University of Texas at Austin for providing HPC resources that have contributed to the research results reported within this paper. URL: http://www.tacc.utexas.edu.

